# The GTPase-activating protein CG42795 is a potent neuronal regulator of ageing in *Drosophila melanogaster*

**DOI:** 10.1101/2024.06.17.598893

**Authors:** Gergő Falcsik, Fanni Keresztes, Annamária Juhász, Zsófia Simon-Vecsei, Orsolya Kolacsek, Győző Szenci, Szabolcs Takáts, Péter Lőrincz, Tamás I. Orbán, Tibor Kovács

## Abstract

Macroautophagy (hereafter referred to as autophagy) is an important self-renewal process in our cells, whereby potentially harmful cellular components are encapsulated within a double-membrane autophagosome and subsequently fused with a lysosome for degradation by acidic hydrolases. This lysosomal degradation process is essential for maintaining cellular homeostasis. Dysfunctional autophagy can lead to the accumulation of cytotoxic protein forms that contribute to the onset of age-related diseases. However, it has been proven that autophagic activity declines with age, so it is therefore particularly important to stimulate autophagy during the lifespan of post-mitotic cells, such as neurons, where cell division is not a possibility in order to replace dead cells. Our research group aims to find new autophagy activation sites to stimulate the efficiency of acidic degradation in neurons during ageing. One approach is the stimulation of membrane fusion events, which are necessary for autophagic degradation, through the activation of small GTPase enzymes. Our previous results have shown that neuron-specific overexpression of the activated form of the Rab2 small GTPase has autophagy and lifespan-enhancing effects. In the present study, we used an RNA interference screen to investigate whether silencing 12 GTPase-activating proteins (GAPs) belonging to the TBC1 domain family can enhance Rab2 activation in the *Drosophila* nervous system. Several of the GAPs studied increased the number of Rab2-positive structures, 5 of which were selected for further screening. Our results suggest that neuronal silencing of *CG42795* exerts an effect on autophagy, with the capacity to enhance the locomotor ability of animals and prolong lifespan. Furthermore, the human orthologue of this GAP, the TBC1D30 protein showed a conserved function in HeLa cells. Silencing TBC1D30 increased the number of active Rab2 protein and enhanced autophagic activity in human cells. Our findings suggest that studying GAPs could be a promising new field of focus for ageing researchers. Further analysis of CG42795 and TBC1D30 could lead to the development of potential autophagy activators.

## Introduction

Cells utilise degradation processes called autophagy or cellular self-digestion to remove damaged organelles and protein aggregates, as well as to create new building blocks. The intensity of autophagy must be tightly regulated to ensure cellular homeostasis [1], [2]. Autophagy comprises three distinct processes: microautophagy, chaperone-mediated autophagy (CMA) and macroautophagy, each of which plays a critical yet discrete role. Microautophagy directly engulfs cytoplasmic components by the invagination of the lysosomal membrane [3]. Conversely, CMA is capable of degrading soluble proteins that are selectively and directly translocated to the lysosome. All substrate proteins contain a KFERQ-like motif that is recognized by a chaperone complex (containing hsc70) in the cytoplasm. It has been observed that this complex interacts with the lysosomal associated membrane protein type 2A (LAMP-2A) to assist the translocation step [4]. Macroautophagy (hereafter autophagy) is a type of catabolic degradation in which cytoplasmic compartments are sequestered within a double-membrane vesicle, the autophagosome, which fuses with lysosomes to deliver the degradation enzymes and create the environment in which the enzymes can function and eliminate the particles [5].

The precise location of phagophore formation remains to be elucidated, although the endoplasmic reticulum (ER) has been identified as a primary regulator. However, mitochondrial, Golgi and plasma membrane origins have also been described [6], [7]. Phagophore formation is initiated by the activation of the Atg1 (autophagy-related 1; ULK in mammals) kinase complex, which phosphorylates and activates other proteins required for autophagosome biogenesis [8]. The TOR (target of rapamycin; TORC1 in mammals) pathway, which is the master regulator of intracellular metabolism, is responsible for regulating the Atg1 kinase complex. If the pathway detects elevated levels of certain nutrients, amino acids or insulin, it blocks autophagy by inhibiting the Atg1 (autophagy-related 1) kinase complex, since the cell has sufficient building blocks and energy for protein synthesis. However, in conditions of nutrient deprivation, the TOR signaling pathway is deactivated, resulting in the activation of the Atg1 kinase complex and the formation of new autophagosomes. This process enables the cell to obtain building blocks from the breakdown of its own materials, because protein synthesis must be continuous in a cell, even in the presence of nutrient deficiency. This regulatory mechanism underscores the critical role of autophagy in cell survival, beyond its mere function in recycling [9].

The initial stages of autophagosome biogenesis also involve the Vps34/PI3K complex, which is responsible for the conversion of phosphatidylinositol (PI) to phosphatidylinositol 3-phosphate (PI3P) on the phagophore membrane. PI3P, a specific PI of the autophagosome membrane, is recognized by other proteins essential for autophagy, leading to their recruitment to the forming membrane [10], [11]. One such example is Atg2, which forms a complex with Atg18. The Atg2-Atg18 complex channels lipids into the phagophore membrane, increasing its surface area. The membrane also incorporates small vesicles containing Atg9 transmembrane proteins, a process that contributes to its growth. In addition to PI3P, another important identity-determining element is added to the membrane of the phagophore and subsequently the autophagosome, namely the Atg8 (LC3B in mammals, Atg8a in *Drosophila*), a ubiquitin-like protein. The ubiquitin ligase-like conjugation system facilitates the attachment of phosphatidylethanolamine lipid anchor to the free Atg8 proteins. These lipidated Atg8 proteins are then anchored to the phagophore membrane. Amino acid cleavage of the Atg4 protein at the C-terminus of Atg8 is required for lipid anchor placement. Atg8 has the capacity to interact with the adapter protein p62 (*Drosophila* ortholog Ref(2)P). It has been observed that many substances marked for selective degradation are processed by Atg8-p62 (and other adaptors that also linked to Atg8) proteins [12]. Researchers often use mutant, fluorescent forms of these two proteins as autophagy markers.

The formation, transport, and fusion of vesicles during the degradation pathway are meticulously regulated processes that require a multitude of proteins with distinct functions. These include small GTPase, coat, motor, docking and folding proteins, among numerous others [8].

Optimal levels of autophagy are required for cell survival, and impairment of autophagy is associated with age-related diseases such as neurodegeneration and cancer [13], [14]. Autophagic activity has been shown to decrease over the lifespan, contributing to the accumulation of toxic protein aggregates within cells, thereby resulting in the loss of functionality and viability [15], [16], [17], [18]. This decline is particularly problematic in post-mitotic cells, such as neurons, where the loss of damaged neurons is irreversible [19]. Moreover, studies have revealed that the neuronal autophagic process is generally suppressed in its downstream stages, leading to the accumulation of undigested particles in autolysosomes [20].

As autophagy gradually declines with age, resulting in neuronal death and neurodegenerative diseases, it is necessary to identify novel targets to enhance autophagy. A promising avenue for enhancing autophagy’s efficacy involves the promotion of lysosomal fusion events, a process that is often compromised with age. The HOPS and SNARE complexes regulate the fusion steps with the assistance of small GTPase proteins [21]. Our previous studies have focused on three small GTPase proteins: Rab2, Rab7 and Arl8, all of which have important and evolutionarily conserved roles in degradation. Of these Rab2 has been shown to induce not only autophagosome-lysosome but also Golgi vesicle-lysosome fusion steps. The function of small GTPase proteins includes membrane fusion, tracking, recruitment of proteins and membrane docking [22]. Small GTPases act like molecular switches, alternating between active GTP-bound and inactive GDP-bound forms [23]. There are two types of enzymes that regulate small GTPases: GAP enzymes (GTPase-activating proteins) induce GTP hydrolysis and inactivate the protein, while GEFs (guanine nucleotide exchange factors) promote GTP binding and activation [24].

Our previous results have demonstrated that Rab2 activation exerts a positive effect on lifespan, movement and autophagic activity. Constitutively active Rab2 has also been shown to be beneficial in the *Drosophila* model of Parkinson’s disease, where it has been observed to extend lifespan, improve degradation and reduce the amount of toxic alpha-synuclein particles in the brain [25].

The present investigation is focused on the identification of GAP (GTPase-activating protein) enzymes that modulate Rab2 activity. By selectively targeting specific GAPs, it is possible to interfere with the inactivating enzymes involved in this process, thereby activating the small GTPase protein. It is hypothesized that this will have a similar effect to constitutively active Rab2 and increase autophagy, lifespan and locomotion. An RNA interference (RNAi) screen was used to investigate 12 GAPs belonging to the TBC1 domain family to find Rab2 regulators.

## Materials and Methodes

### Strains used for experiments and their maintenance

We used *Drosophila melanogaster* for our experiments. The strains have been obtained from the Bloomington Drosophila Stock Centre (BDSC) and have also been generated by crossing animals from the ELTE Department of Genetics Strain Collection (EDSC) **(*Table 1*)**. The strains used were maintained on a standard corn-sugar-agar medium. All our experiments were carried out on animals kept at 29°C. The 12 GAP genes silenced in the experiment are referred to by their CG number or as GAP-RNAi. For control animals we used an isogenic *w[1118]* (*isowhite*) strain simulating the wild-type *Drosophila*, and for certain experiments an additional RNAi-control strain (GFP-RNAi) were used **(*Table 1*)**. This may be necessary because the double-stranded RNA expressed in RNAi strains can itself induce non-specific effects by activating the RISC complex. The RNAi control is designed to simulate the non-specific effects of RNAi by producing extra targetless double-stranded RNA compared to the *isowhite* (wild type with white eyes) control [26]. However, this GFP-RNAi control strain cannot be used in experiments where the expression of the GFP protein is required (for example, in the case of several of our fluorescent microscopy experiments).

**Table 1:** Name, genotype, and place of origin of the strains used in our experiments.

| Name | Genotype | Origin |
| --- | --- | --- |
| control / isogenic white | w[1118] | BDSC: 5905 |
| GFP-RNAi (RNAi-control) | y[1] sc[*] v[1]; P{y[+t7.7] v[+t1.8]<br>=VALIUM22-EGFP.shRNA.4}attP2 | BDSC: 41551 |
| CG1695 (GAP-RNAi) | y[1] v[1]; P{y[+t7.7] v[+t1.8]<br>=TRiP.HMJ24138}attP40 | BDSC: 62898 |
| CG4041 II (GAP-RNAi) | y[1] sc[*] v[1] sev[21]; P{y[+t7.7] v[+t1.8] =TRiP.GL00239}attP2/TM3, Sb[1] | BDSC: 35332 |
| CG4041 III (GAP-RNAi) | y[1] sc[*] v[1] sev[21]; P{y[+t7.7] v[+t1.8] =TRiP.GL00239}attP2/TM3, Sb[1] | BDSC: 35332 |
| CG5344 (GAP-RNAi) | y[1] sc[*] v[1] sev[21]; P{y[+t7.7] v[+t1.8] =TRiP.HMC06579}attP40 | BDSC: 77447 |
| CG6182 (GAP-RNAi) | y[1] sc[*] v[1] sev[21]; P{y[+t7.7] v[+t1.8] =TRiP.HMC06532}attP40 | BDSC: 77400 |
| CG7112 (GAP-RNAi) | y[1] sc[*] v[1] sev[21]; P{y[+t7.7] v[+t1.8] =TRiP.HMS01132}attP2/TM3, Sb[1] | BDSC: 34976 |
| CG7324 (GAP-RNAi) | y[1] sc[*] v[1] sev[21]; P{y[+t7.7] v[+t1.8] =TRiP.HMS00722}attP2 | BDSC: 32929 |
| CG8085 (GAP-RNAi) | y[1] v[1]; P{y[+t7.7] v[+t1.8] =TRiP.JF03085}attP2 | BDSC: 28670 |
| CG11727 (GAP-RNAi) | y[1] v[1]; P{y[+t7.7] v[+t1.8] =TRiP.HMS01818}attP40 | BDSC: 38350 |
| CG12241 (GAP-RNAi) | y[1] sc[*] v[1] sev[21]; P{y[+t7.7] v[+t1.8] =TRiP.HMS00612}attP2/TM3, Sb[1] | BDSC: 33729 |
| CG31935 (GAP-RNAi) | y[1] sc[*] v[1] sev[21]; P{y[+t7.7] v[+t1.8] =TRiP.HMC05513}attP40 | BDSC: 64495 |
| CG42612 (GAP-RNAi) | y[1] sc[*] v[1] sev[21]; P{y[+t7.7] v[+t1.8] =TRiP.HMC06174}attP40 | BDSC: 66313 |
| CG42795 (GAP-RNAi) | y[1] v[1]; P{y[+t7.7] v[+t1.8] =TRiP.HMJ23781}attP40/CyO | BDSC: 62385 |
| DdcGal4 | w[1118]; P{Ddc-GAL4.L}4.36 | BDSC: 7009 |
| DdcGal4-YFP-Rab2 | w[1118]; DdcGal4; YFP-Rab2/ST6 | EDSC |
| cgGal4 (II.) | w[1118]; P{Cg-GAL4.A}2 | EDSC |
| cgGal4-YFP-Rab2 | w[1118]; cgGal4; YFP-Rab2/ST6 | created by us |
| YFP-Rab2 | y[1] w[*]; P{w[+mC]=UAST-YFP.Rab2}l(3)neo38[02]/TM3, Sb[1] | BDSC |
| Sp;Sb/ST6 | w[*]; Sp; Sb / ST6 | EDSC |
| 3x mCh-Atg8a, DdcGal4 | w[1118]; DdcGal4; 3x mCh-Atg8a | EDSC |
| Act<CD2<Gal4,GFP | hsFlp; pAct<CD2<Gal4 UAS-nlsGFP | gift from Gábor Juhász |
| gln-GFP-Rab2 | w[*]; gln-GFP-Rab2 | gift from Ole Kjaerulff |
| CG42795Mi | y[1] w[*]; Mi{y[+mDint2]=MIC}CG42795[MI11017] | BDSC: 55572 |

For clonal systems, we used the strain Act<CD2<Gal4,GFP (V40) ***(Table 1)***. The essence of this strain is that between the coding sequence of the Gal4 protein and the actin promoter that drives it, there is a flip out cassette that prevents transcription of the gene. There are FRT sequences at both ends of the flip out cassette where the FLP protein can loop out this sequence, allowing the Gal4 protein to be transcribed again. The FLP protein can be activated by heat shock (hsFLP) in random cells. Where the FLP has been activated by heat shock and the Gal4 protein can be transcribed, these cells express GFP as well as the GAP-RNAi sequence, but in the other cells the transgenes are not expressed in the absence of Gal4, these cells can be considered as “wild type” control cells [27].

### Fluorescence microscopy

We studied the effects of GAP-RNA interference in two types of tissues, adult *Drosophila* brains and larval fat bodies. Adult female flies were decapitated after anaesthetized with ether, the heads were dissected and fixed in formaldehyde (4%) diluted in PBS (phosphate-buffered saline) for 20 min at room temperature, then washed 3 x 5 times in PBS. We then dissected the brains, also in PBS, and covered the samples with Hoechst solution containing PBS:glycerol (ratio 1:4) before microscopy. To examine fat bodies, the fat body lobes of larvae at the L3 feeding stage (L3F) were dissected in PBS without fixation. Each sample was made from approximately 5-6 larvae. After dissection the samples were immediately covered with Hoechst solution containing PBS:glycerol (ratio 1:4). For starvation experiments, L3 feeding stage (L3F) larvae were transferred to a 20% sucrose solution (protein starvation) for 2 hours prior to dissection. Then they were dissected in PBS without fixation and covered with Hoechst solution containing PBS:glycerol (ratio 1:4).

Microscopic images were taken using a Zeiss Axioimager Z1 microscope equipped with ApoTome2. The brains were photographed at 200x magnification, while the fat bodies were photographed at 400x magnification. The images were taken across the entire thickness of the organ, at a distance of 1 µm per slice. During data analysis, we merged the slices of the organ so that all the fluorescently labeled structures could be counted. We used ZEN 3.5 (Blue edition v3.5.093) and ImageJ software to analyze the images.

In the experiments with the clonal systems, the larvae were transferred to 29°C for 18 hours prior to dissection, and except for this period, the larvae were kept at 25°C. For LysoTracker Red (L7528) staining of acidic compartments in fat bodies, we diluted LysoTracker Red 1:1000 in PBS. After staining for 2 minutes, we washed the samples twice in PBS for 2-2 minutes before covering. The samples were covered with Hoechst solution in PBS:glycerol (ratio 1:4).

### TBC1D30 knock down

HeLa cells were cultured in Dulbecco’s modified Eagle’s medium (DMEM) supplemented with 10% fetal calf serum, 1%L-glutamine and 1% penicillin/streptomycin (Gibco). Prior to siRNA transfection 0,5x105 cells were seeded onto a 12 well plate. Next day 30 pmol Silencer Pre-designed siRNA specific against human TBC1D30 (Invitrogen, AM16708 ID: 264758) was transfected with Lipofectamine RNAiMAX reagent (Invitrogen) by the manufacturer’s instructions. For negative control experiment 30 pmol Silencer Select Negative Control No.1 siRNA (Invitrogen, 4390843) was transfected in parallel. Knock down efficiency was controlled in three independent experiments at 48h posttransfection by RT-qPCR. Harvested cells were lised in TRIzol Reagent (Invitrogen), total RNA was isolated and reverse transcribed with High-Capacity cDNA Reverse Transcription Kit (Applied Biosystems) according to the manufacturers instructions. 5 μl ten-fold diluted cDNAs were used in 20 μl PCR reaction volume in triplicate. TBC1D30 TaqMan assay (Applied Biosystems, Hs00392437_m1) in TaqMan Gene Expression Master Mix (Applied Biosystems) was used for measurements in QuantStudio 3 platform with the following thermal profile: 95 °C 10 min, 40 cycles of 95 °C 15 s and 60 °C 1 min. POLR2A TaqMan assay (Applied Biosystems, Hs00172187_m1) was used for endogenous control. Quantification was carried out by the ΔΔCt method with Diomni Design and Analysis (RUO) 3.1.0 software. (Error bars represent 95% confidence intervals.)

Knock down fluorescent reporter HeLa cell line was analysed at 72h posttransfection by confocal microscopy. For colocalisation experiment and immunfluorescent staining transfected HeLa cells were harvested at 48h posttransfection and 1,5x104 cells were seeded onto a 0,8 cm2 chamber. Next day cells were washed with cold PBS and fixed in 4% PFA for 15 min at 4°C. Cells were then washed three times with PBS and stored at 4°C for subsequent immunostaining and confocal microscopy.

### Drosophila Immunohistochemistry and Colocalisation experiment in human cells

Immunocytichemistry experiment was performed as earlier [28] with the following modifications. Anti-Rab2A (Proteintech, produced in mouse) and anti-TBC1D30 (ThermoFisher Scientific, produced in rabbit) antibodies were used as primary antibodies, Alexa568 conjugated anti-mouse (Invitrogen) and DyLight488 conjugated anti-rabbit (Invitrogen) antibodies were used as secondary antibodies. The doubly labeled cells were examined with a confocal microscope, during which thin (1µm) slices were recorded along the Z-axis of the cells. At the evaluation of the obtained confocal microscopy images, the cell nucleus was excised and omitted from the analysis, as it showed high autofluorescence. We calculated a Pearson correlation based on the microscopic images, which measures the strength of the linear relationship between the green and red channel intensities at pixel level. The axes show the strength of each fluorophore: weak signal intensity is near the origin, and strong signal intensity is further away. The stronger the Pearson correlation, the more the two fluorophores are arranged along a slanted line in the samples. The Pearson coefficient, also known as Pearson’s correlation coefficient, is a statistical measure that indicates the strength and direction of a linear relationship between two variables. In our experiment, the Pearson r value indicates whether the intensity of the green channel increases in line with that of the red channel from pixel to pixel.

### Drosophila Immunohistochemistry

We used immunohistochemistry to study the quantitative changes in ubiquitinated structures that are present in the brain of the fruit flies. For the experiment, we used the brains of 7-day-old female *Drosophila*. This was done as described by Kovács et al [20]. The samples were prepared as described for fluorescent microscopy and fixed in formaldehyde (4%) for 40 minutes at room temperature. After fixation, we did the washing with PBT (3x 10 minutes, PBS + 0.1% Tween20). Then, we dissected the brains and blocked them for 1 hour in 1ml PBT + 50μl FBS (fetal bovine serum) solution. The samples were then incubated with the primary antibody: anti-ubiquitin (mouse, 1:500, Merck, ST1200) at 5°C for 2 days. Excess primary antibody was washed out with PBT (3x 20 minutes), followed by another blocking for 20 minutes, after which we incubated the brains with the secondary antibody: Alexa Fluor 488 anti-mouse (1:500, Life Technologies, A11001) for 2 hours at room temperature. We also washed out the excess secondary antibody in several steps as described for the primary antibody, and then, during post-dissection, we removed the remaining tissue from other organs and tracheas from the surface of the brains before microscopy. The nuclei were labeled with Hoechst containing PBS:glycerol (1:4) nuclei stain.

### Western blot

To measure quantitative changes in certain proteins, we used western blotting. For the measurement, we isolated protein from the heads of 7-day-old female animals. The animals were anesthetized with ether and then decapitated. We collected 15-15 heads from each strain in Eppendorf tubes. Proteins were extracted from the heads in a 1:1 mixture of 96 μl Fly Lysis Buffer and 2xLaemmli buffer (Bio-Rad, 1610737) using a homogenizing rod and then stored at -20°C. We performed the Western blot according to the method described by Billes et al [29]. We applied 15μl of each sample. The samples were run on a 4-20% Mini-PROTEAN® TGX™ gradient gel and then blotted onto a nitrocellulose membrane (Kisker Biotech, 40520100). After that, we blocked the samples in 3% milk powder solution (BioRad 170–6404 Blotting-Grade Blocker) for 1 hour, and then applied the primary antibodies: anti-Ref(2)P/p62 (rabbit, 1:2000 [30]), α-Tub84B (internal control) (mouse, 1:1000, Sigma, T6199), anti-Atg8a (rabbit, 1:2000 [31], [32]). We incubated the sample in the antibody solutions at 5°C overnight. The next day, the excess (unbound) primary antibody was washed off the nitrocellulose with TBST, 3 x 10 min washes. Then we incubated the nitrocellulose in the secondary antibodies: anti-rabbit IgG alkaline phosphatase (1:1000, Sigma, A3687), anti-mouse IgG alkaline phosphatase (1:1000, Sigma, A5153) for one hour. Detection was done with NBT-BCIP solution (1:50, Sigma, 72091). We used the ImageJ software to evaluate the results. The levels of p62 and Atg8a were corrected by the change measured for α-Tub84B. Western blotting is a semi-quantitative technique. Therefore, we only recorded the direction of changes, not their extent.

### Lifespan analysis

The goal of this experiment is to compare the average lifespan of each gene silenced strain to the wild type animals. The measurement was performed with 5 parallel tubes, 15 female and 15 male animals of each genotype were placed in vials. Every two days, we transferred the *Drosophila* to fresh feeding vials. The number and sex of the deceased fruit flies was recorded on a daily basis until the complete mortality of the sample was achieved.

### Climbing test

In this experiment, the hypothesis can be tested by comparing the fitness and state of the nervous system of the fruit flies as they age. In a small-diameter, long glass tube, Drosophila start to climb upwards after being knocked down (negative geotaxis). The speed of perception and retention of the stimulus is proportional to the state of the fly’s nervous system. The short-term climbing test measures the ability and speed of the animals to perceive the stimuli [33]. We measured the climbing ability of fruit flies at 7, 14, 21 and 28 days of age. Measurements were made with two parallel tubes. We anesthetized each genotype with CO2, then separated the sexes and placed 10-10 animals in each tube. The animals were then rested at 29°C for 1 hour. After resting, the climbing test was repeated three times. There was a 30-minute break between each test. Each trial was recorded on video file and used to evaluate how each genotype was performing. Short-term climbing takes place to the first boundary line (6 cm) and measures how many of 10 animals reach the boundary line within 20 seconds.

### Statistics and data representation

We used RStudio (2021.09.0 Build 351) to perform the statistical analysis of the experimental data. The first step was to test the distribution of the data. In the case of a normal distribution, we used the t-test, in the case of a non-normal distribution, we used the Mann-Whitney U-test. For the results of the clonal systems, we performed the statistics with a paired t-test in the case of a normal distribution (Two Sample t-test if the two data set has similar variance, Welch Two Sample t-test if the two data set didn’t have similar variance), and a paired Mann-Whitney U-test in the case of a non-normal distribution. The confidence level of significance is indicated by the stars on the graphs: **p<0,05:*, p<0,01:**, p<0,001:\*\*\***. Black stars indicate significance compared to wild type (*isowhite*) control; brown stars indicate significance compared to RNAi control (*GFP-RNAi*).

We created box plots using the RStudio program to represent the evaluated data. In the box plot, the box represents the values belonging to the interquartile range (50% of the data), and the black line in the boxes represents the median. The lower horizontal line shows the minimum value, and the upper horizontal line shows the maximum value. Outliers in the data set are indicated by empty circles. The black dots on the graph represent a value for a single sample, e.g. the evaluation of a microscopic image, except for the climbing essay, where one dot represents a measurement, e.g. the climbing ability of 10 animals in percentages. In the case of box plots, the data have been normalized. We generated the Kaplan-Meier curves of lifespan assay measurements and analyzed data using SPSS22.0 software. The data type of the lifespan assay cannot be displayed using boxplot, so we used a bar chart instead. The standard deviation is shown in the tracks at the top of the bars.

## Results

### The depletion of specific GAPs results in increased levels of active Rab2 protein

The present section of our research is oriented towards the investigation of the effect of silencing 12 GAP proteins potentially involved in the regulation of Rab2 and thus in the network regulating autophagic-degradation pathways ***(Figure 1)***.

**Figure 1:**
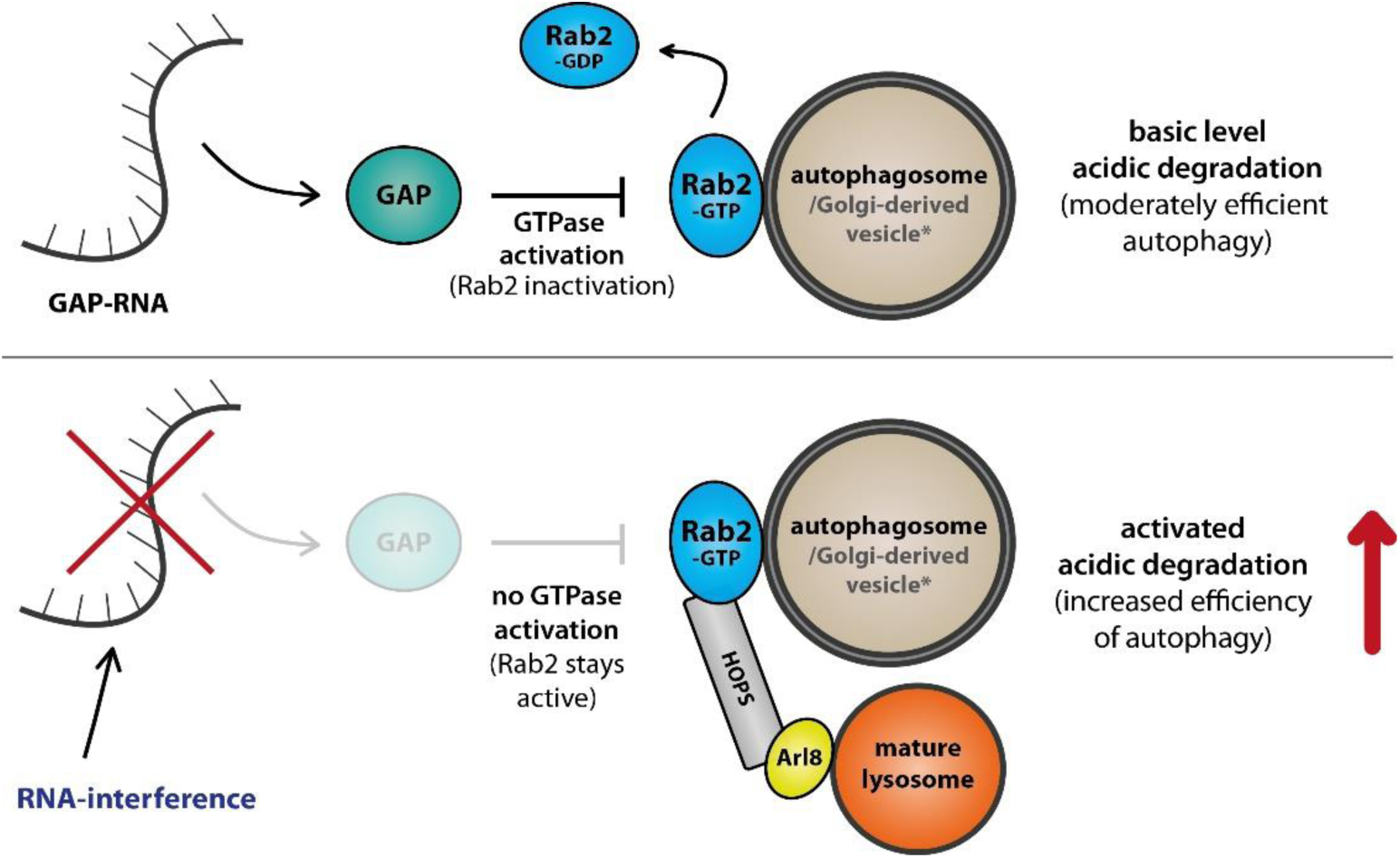
GAP screen. Enhancing autophagic degradation efficiency through silencing regulatory GAPs. Rab2-specific GAP activates the hydrolysis of GTP, thereby converting membrane-bound, active Rab2 proteins into an inactive, soluble form (top figure). By silencing the right GAP, Rab2 remains active and the efficiency of autophagic degradation pathways can be increased (bottom figure). *In several organs, fusion of the autophagosome and vesicles containing Golgi-derived hydrolase enzymes with the lysosome are both linked to Rab2. Stimulation of both fusion events can increase degradation efficiency.

In our initial experiments, we investigated which GAP gene silencing could increase the amount of active Rab2 protein — important for vesicle fusion — in larval fat body and nervous system of *Drosophila melanogaster*. We started with the fat body screen because the *Drosophila* larval fat body is a preferred tissue for studying autophagy, since their cells are large and show changes in autophagic structures well [34]. For this screen, we crossed the RNAi strains with *cgGal4; YFP-Rab2* flies. The cgGal4 (collagen promoter-Gal4) is specifically expressed in the larval fat body [35]. We performed the assay in two conditions, using well-fed and starved animals. As a result of starvation, autophagy is activated in the animals, while under well-fed conditions, few autophagic vesicles can be detected [36]. Of the 12 genes 4 GAP-RNAi (*CG6182*, *CG8085*, *CG31935* and *CG42612*) were effective in well-fed larvae ***(Figure 2: A-B)***, and in starving larvae 6 (*CG1695, CG8085, CG11727, CG12241, CG31935*, *CG42795*) ***(Figure 2: B)*** of them were able to significantly increase the amount of active Rab2 proteins. Some interesting patterns emerged in the well-fed larvae when the GAP silenced strains were examined. While there were hardly any Rab2-positive vesicles in the control strain, many small structures appeared after *CG8085* was silenced ***(see Figure 2: A’)***. In the *CG31935*-RNAi strain, fewer but larger vesicles were visible, mainly at the periphery of the cells ***(Figure 2: A”)***. Also in fat-bodies, we generated clone cells by somatic recombination using the Act<CD2<Gal4,GFP system ***(Table 1)***. Because of the mechanism described in the materials and methods section, the Gal4 protein sequence driving the GFP and RNA-interference genes is expressed only in clone cells in these animals. Surrounding cells in which the Gal4 protein is not transcribed are considered control cells. The amount of acidic degradative vesicles was assessed using LysoTracker-Red staining. LysoTracker-Red marks such acidic organelles as lysosomes, autolysosomes and late endosomes. As with the Rab2 screen in fat bodies, we also tested the larvae here in both well-fed and starving conditions. In well-fed animals, silencing of 4 GAP genes (*CG1695*, *CG7324*, *CG11727*, *CG42612*) increased the amount of acidic vesicles in the clone cells ***(Figure 2: C)***. For starving larvae 2 GAP silencing (*CG11727* and *CG31935*) increased, 4 silencing (*CG4041(II)*, *CG4041(III)*, *CG6182*, *CG7324*) decreased the number of acidic vesicles in clone cells compared to control cells ***(Figure 2: D)***. In some cases, there was a large increase in the number of acidic vesicles ***(Figure 2: D)***, which may indicate that the degradative processes were upregulated. This can be beneficial for breaking down harmful structures in these cells.

**Figure 2:**
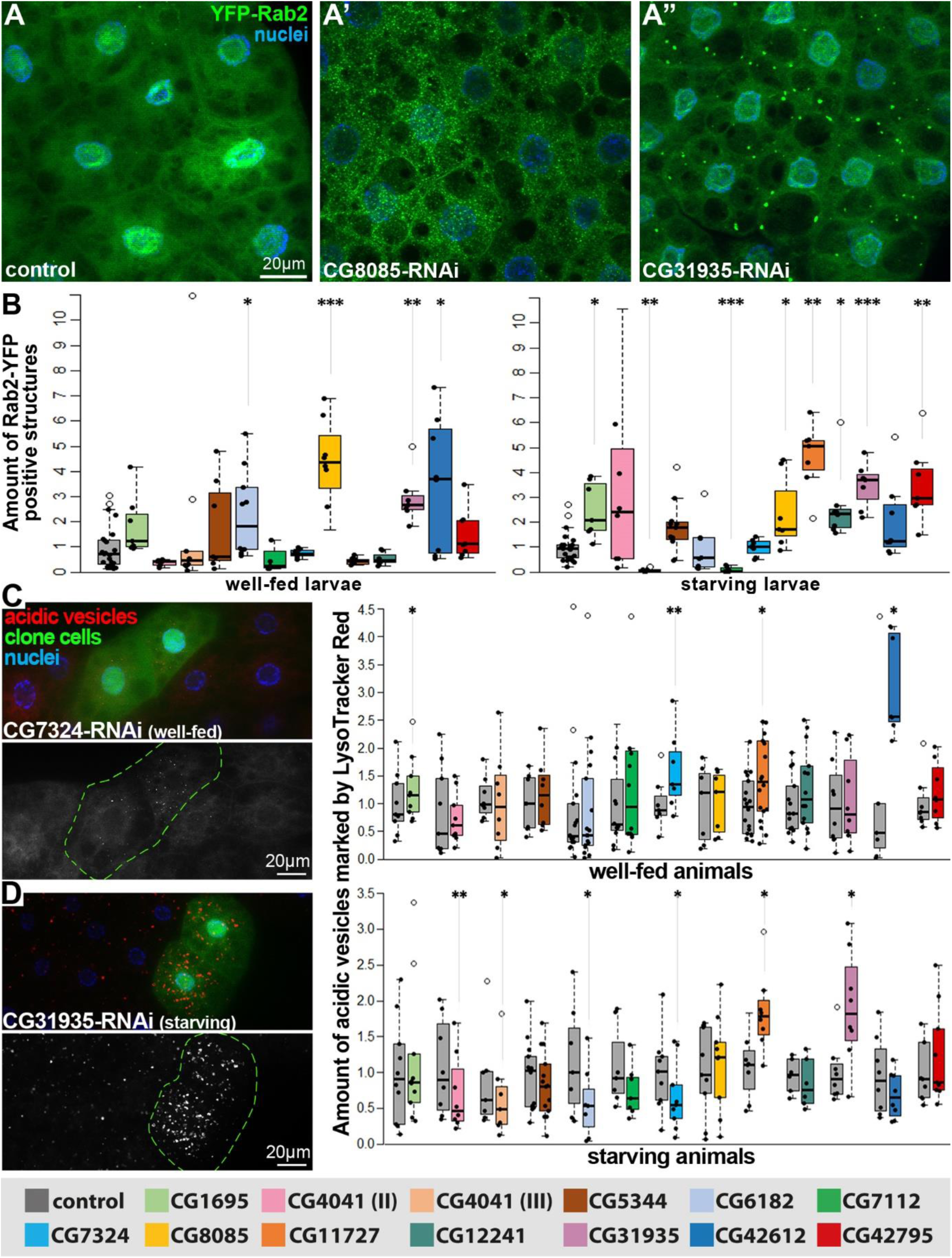
GAP silencing can increase the number of Rab2 positive structures in Drosophila brain and larval fat body. A-A”: Images of changes in the amount of Rab2-positive structures in larval fat cells due to certain GAP silencing. All three fluorescent images captured from well-fed larvae. **B:** Diagrams showing the evaluated data measured in from larval fat bodies after GAP silencing. In well-fed larvae: left, and in starving larvae: right plot. **C:** Clonal cells expressing GFP and RNA, silencing the gene of interest in the fat body of well-fed larvae, on the left. And the evaluated data on the right. **D:** Clonal cells showing a significant increase in acidic vesicles (red spots). The evaluated data is on the right. **Levels of significance: * = p<0.05; ** = p<0.01; *** = p<0.001**

### Neural autophagy can be activated by suppression of GAPs

During neuronal screen, the individual RNA interference lines were crossed with *DdcGal4; YFP-Rab2* animals. This tissue-specific Dopa decarboxylase (Ddc) promoter only allows Gal4 expression in the dopaminergic and serotonergic neurons of the brain [37]. Our focus was on the mushroom body (MB) region of the brain, which contains a large number of dopaminergic and serotonergic neurons and is suitable for modeling neurodegenerative diseases of old age [38], [39], [40]. We examined the cell body and axonal regions of neurons separately because neurons are highly polarized cells and there can be large differences between the two cell regions ***(Figure 3: A-B’)***. Of the 12 GAP proteins, 7 GAP silencing (*CG1695*, *CG5344*, *CG7324*, *CG11727*, *CG12241*, *CG31935* and *CG42795*) were able to increase the amount of active Rab2 protein at the cells axonal area, and 6 of them (*CG1695*, *CG5344*, *CG11727*, *CG12241*, *CG31935* and *CG42795*) at the perikaryon ***(Figure 3: C)***. These results of the two RNAi screen suggest that these genes may be regulators of Rab2 protein activity, either directly or indirectly, in *Drosophila* neurons and larval fat bodies. Thereby their downregulation could be a cause of the increase of the Rab2 protein active form. Disabling these GAP genes might lead to enhanced autophagic-degradation in cells.

**Figure 3:**
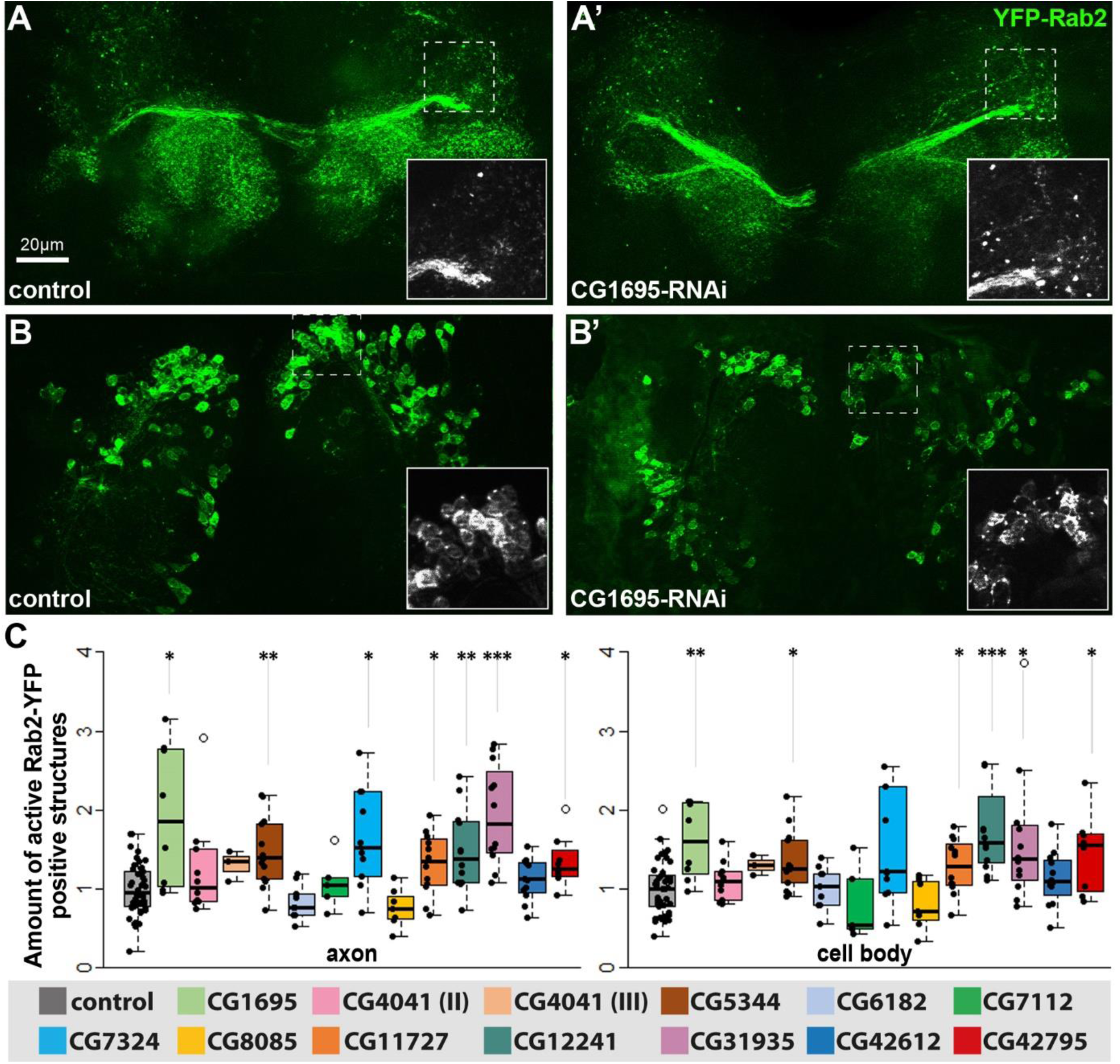
GAP silencing can increase the number of Rab2 positive structures in Drosophila brain and larval fat body. A, A’: Fluorescent microscopic images of fruit fly dopaminergic and serotonergic neurons in the mushroom body. The images show changes in the number of YFP-Rab2 positive vesicles (well separated green spots) in the axonal area of the neurons. **B, B’:** Fluorescence microscopy images of changes in the abundance of Rab2-positive structures in the cell body of neurons. **C:** Boxplots depicting changes in the amount of active Rab2 protein in axonal and cell body regions following each gene silencing of GAPs. **Levels of significance: * = p<0.05; ** = p<0.01; *** = p<0.001**

Autophagy activity in the nervous system was assessed by measurement of the mCherry-Atg8a reporter protein (fluorescence microscopy) and the amount of ubiquitinated aggregates by immunohistochemistry. mCherry-Atg8a marks the autophagic vesicles from the forming phagophore to the autolysosome [34]. Atg8a (lipidated form) showed a significant decrease in several cases (*CG1695*, *CG7324*, *CG11727*, *CG31935* and *CG42795*), which may indicate more efficient autophagic-vesicle fusion and degradation in silenced strains. This decrease in lipidated Atg8a is also coincides with our previous observations with Rab2 constitutive activation experiments [25]. But there was an exception, the silencing of *CG12241*, where a significant increase could be seen in contrast to other GAP-RNAi-s ***(Figure 4: A)***. This could mean an upregulation of phagophore formation or a less efficient autophagic-vesicular degradation. Larger protein aggregates and defective organelles that can no longer be degraded by the proteasome are dealt with the macrophagic pathway, so measuring the amount of these structures can also be a good indicator [41]. The amount of ubiquitinated aggregates was significantly reduced in 10 GAP silenced strains (*CG1695*, *CG4041(II)*, *CG5344*, *CG7112*, *CG7324*, *CG8085*, *CG11727*, *CG31935*, *CG42612* and *CG42795*) in *Drosophila* brain ***(Figure 4: B)***. These results could be explained by the increased activity of autophagy, which eliminates the ubiquitinated structures marked for degradation. With Western blot we determined the amount of Atg8a and Ref(2)P (/p62) proteins isolated from the heads of *Drosophila*, and the level of α-Tub84B was also measured as inner control ***(Figure 4: C)***. The ortholog of Atg8a is the human LC3B [33] There are two forms of this protein, the soluble (Atg8a-I) and the lipidated, membrane-bound (Atg8a-II) which is the active form [42]. The level of Atg8a-II (lipidated form) is consistent with the microscopic results, the amount of protein decreases in the RNAi strains. Ref(2)P/p62 protein is degraded during autophagic-lysosomal degradation, therefore it is a good indicator of the efficacy of autophagy [34]. The p62 level is also decreased in all RNAi strains (taking into account the calculation corrected for the amount of α-Tub84B), which according to the evaluation is significant in the case of *CG1695* and *CG42795*, suggesting that autophagy is activated in these cases.

**Figure 4:**
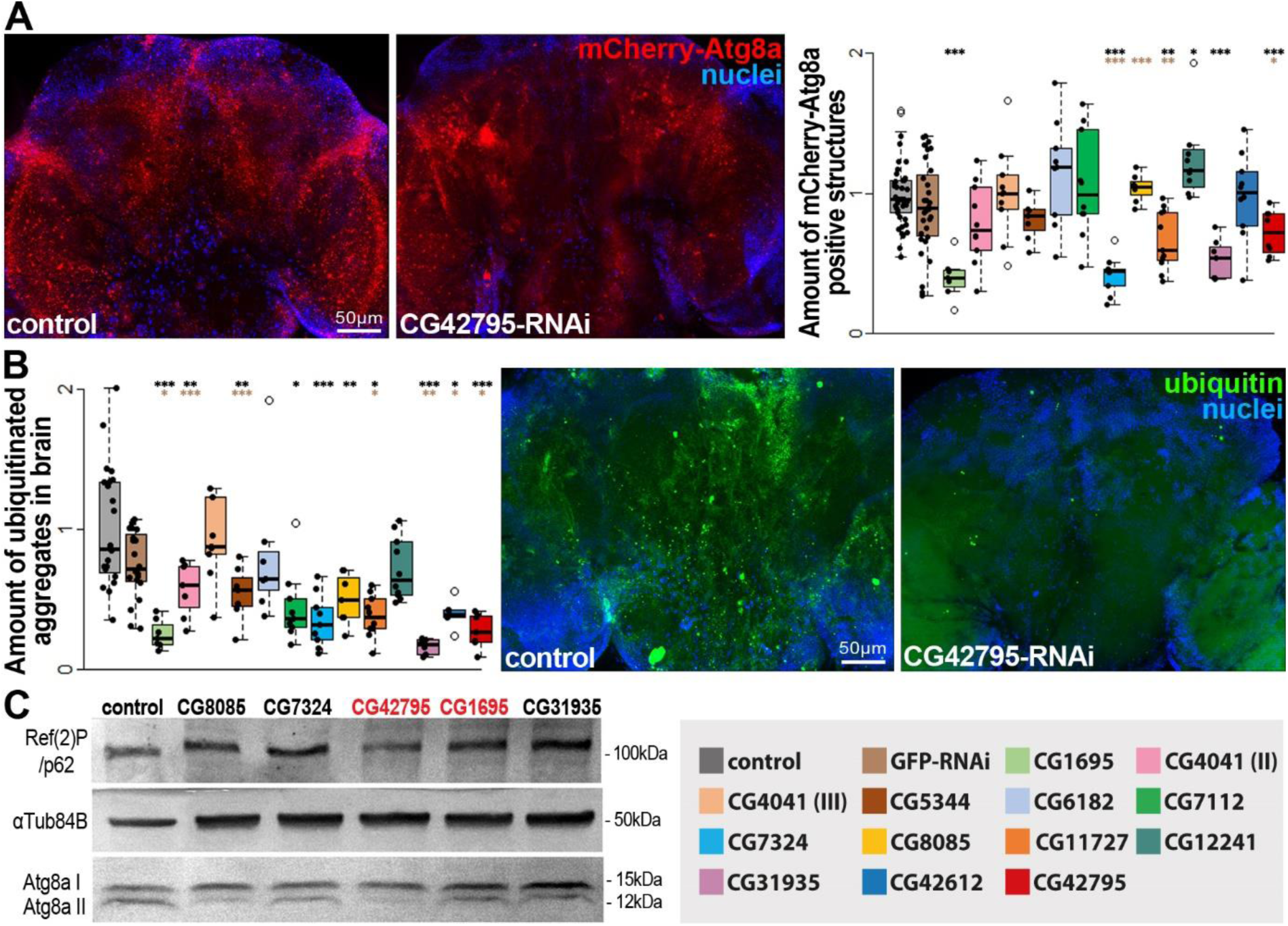
Silencing of certain GAP genes in the Drosophila brain may lead to increased autophagic activity. A: Presence of mCherry-Atg8a (lipidated form) containing autophagic structures in Drosophila brain imaged by fluorescence microscopy. The well separated red spots are the individual autophagic vesicles labeled with the fusion protein marker. And the boxplot representing the evaluated data. **B:** Presence of ubiquitinated aggregates in fruit fly brains labeled by immunohistochemistry after GAP silencing. The boxplot presenting the evaluated data of each silenced strains. Levels of significance: * = p<0.05; ** = p<0.01; *** = p<0.001. Black *: significance compared to wild type animals. Brown *: significance compared to RNAi control animals. **C:** Western blot image of neural protein isolated from 5 chosen GAP silenced strains. Atg8a I = soluble form of Atg8a; Atg8a II = lipidated, bound form of Atg8a.

### Neuron specific silencing of CG42795 and CG8085 extends lifespan in fruit flies

For physiological studies, we selected 5 promising GAP silenced strains (*CG1695*, *CG7324*, *CG8085*, *CG31935* and *CG42795*) that performed well in previous experiments and started with these first. The constitutively active form of the Rab2 protein was able to increase the lifespan of *Drosophila*. If we succeed in silencing GAPs that inactivate Rab2, we can expect a similar result. We also performed the silencing specifically for dopaminergic and serotonergic neurons. Thus, the effects of other tissues or neuron types do not interfere with the experiment. In both sexes *CG8085-* and *CG42795*-RNAi significantly increased the fruit flies average lifespan ***(Figure 5: A, A’)***. Silencing of the other three GAP genes only shows changes in one sex. *CG7324*-RNAi increases the average lifespan of males, but there is no significant change in the lifespan of females. *CG1695*-RNAi shows a significant decrease in males and an increasing trend in females. *CG31935*-RNAi significantly increases lifespan in females but causes no changes in males. These results show that some GAP silencing can increase the average lifespan of *Drosophila*, but for other genes there are differences between sexes. For the 5 GAP silenced strains mentioned above, we also performed a climbing test. This experiment will show the state of the neurons in each of the strains studied at different ages. This is tested by observing how quickly the animals respond to the stimulus and how they can retain it [33]. Some GAP silencing improved, while in one case the climbing ability of the animals was significantly impaired in old age ***(Figure 5: B, B’)***. In females, *CG7324*- and *CG1695*-RNAi significantly increased climbing ability, however in males these only could make an improvement compared to the RNA interference control. In males an additional gene-silencing *CG42795*-RNAi could enhance the climbing ability. Silencing *CG31935* significantly impairs the climbing ability of the flies in both sexes. We can say that silencing of certain GAP genes may be able to increase climbing ability in old animals, but we found large differences between the sexes and some silencing had an opposite negative effect.

**Figure 5:**
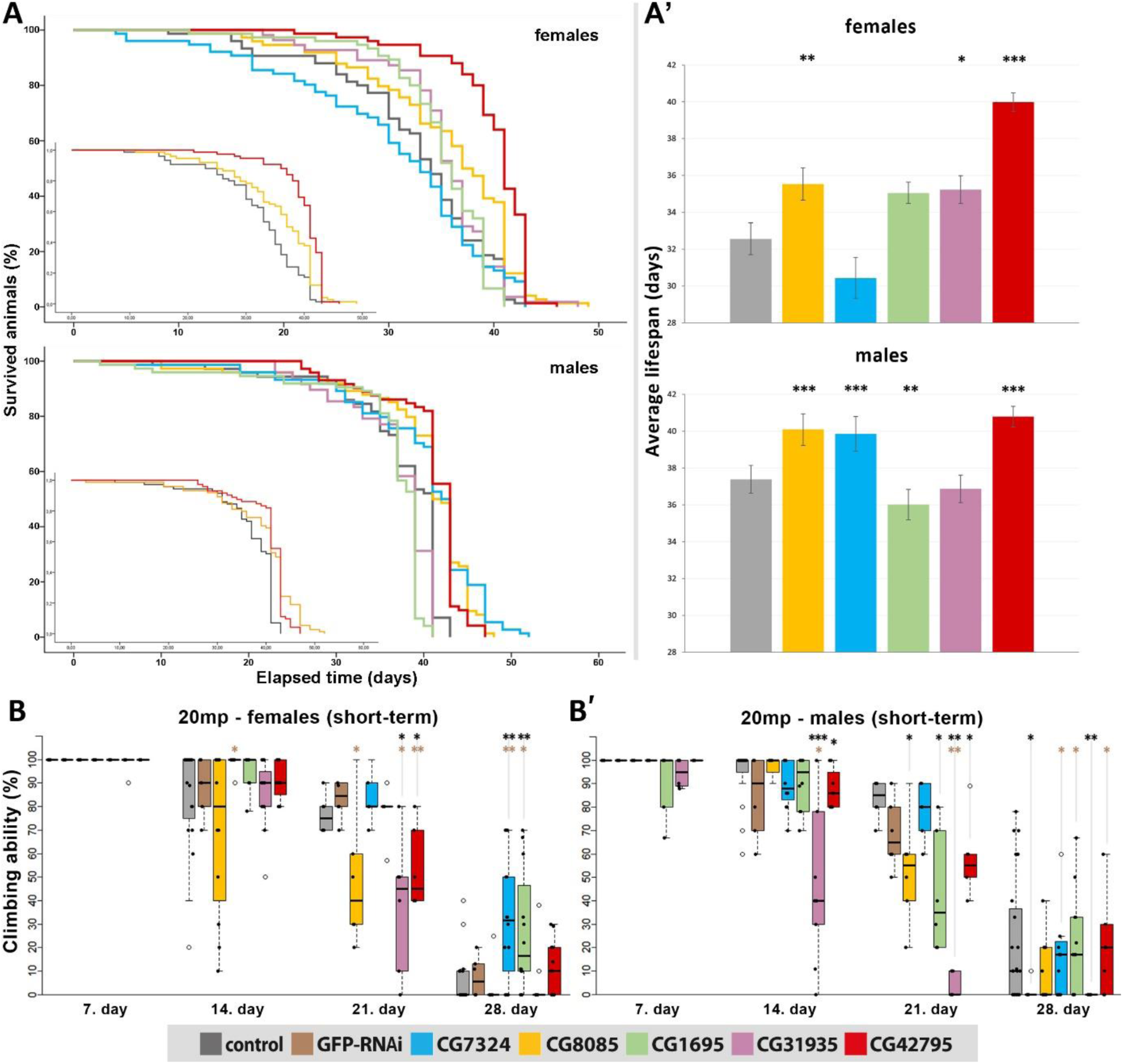
Silencing of CG8085 and CG42795 lead to an increased average lifespan of these Drosophila. A: Kaplan-Meier lifespan curves of individual gene-silenced strains. Curves showing significant increases in both sexes are highlighted by the smaller graphs in the lower left corner. **A’:** Average lifespan plots for each mutant strain, females at top, males at bottom. **B:** Changes in short-term climbing ability of female, GAP silenced animals. **B’:** Changes in short-term climbing ability of male, GAP silenced Drosophila. Levels of significance: * = p<0.05; ** = p<0.01; *** = p<0.001. Black *: significance compared to wild type animals. Brown *: significance compared to RNAi control animals.

We summarized the effects of each GAP silencing in our experiments ***(Table 2)***. The most beneficial effect was observed in experiments with *CG42795* GAP silencing in *Drosophila*. This was able to increase the amount of active Rab2 positive structures in both tissue types studied. It also improved autophagy. Furthermore, it had a positive effect on the average lifespan of the animals and on the climbing ability of aged male animals. *CG1695*-RNAi also performed well, though it has a negative impact on the average lifespan of male flies. Taking these data together we decided to test *CG42795* with a knock-out construct to strengthen our results and in parallel of the mutant experiments we started to investigate the human orthologue of the *Drosophila CG42795*.

**Table 2:** Summary table of the results of GAP screens. From all the investigated GAP-RNAi strains CG42795-RNAi performed the best, showing the most beneficial effects all experiments combined.

| GAP RNAi | Rab2 activation<br>in neurons | Rab2 activation<br>in fat body | autophagic<br>degradation | average<br>lifespan | climbing<br>ability |
| --- | --- | --- | --- | --- | --- |
| CG1695 | ↑ | ↑ | ↑ | ♀⊖ \| ↓♂ | ↑ |
| CG4041 (II) | ⊖ | ⊖ | ↗ | N/A | N/A |
| CG4041 (III) | ⊖ | ↓ | ⊖ | N/A | N/A |
| CG5344 | ↑ | ⊖ | ↑ | ↓ | ⊖ |
| CG6182 | ⊖ | ↑ | ⊖ | ↓ | ⊖ |
| CG7112 | ⊖ | ↓ | ⊖ | N/A | N/A |
| CG7324 | ↑ | ⊖ | ⊖ | ♀⊖ \| ↑♂ | ↑ |
| CG8085 | ⊖ | ↑ | ⊖ | ↑ | ⊖ |
| CG11727 | ↑ | ↑ | ↑ | ↓ | ♀↘ \| ⊖♂ |
| CG12241 | ↑ | ↑ | ⊖ | ♀⊖ \| ↑♂ | ♀↘ \| ⊖♂ |
| CG31935 | ↑ | ↑ | ↗ | ♀↑ \| ⊖♂ | ↓ |
| CG42612 | ⊖ | ↑ | ⊖ | ↓ | ⊖ |
| CG42795 | ↑ | ↑ | ↑ | ↑ | ♀⊖ \| ↑♂ |

= significant increase = increasing trend (non significant) = significant decrease = decreasing trend (non significant) = no change / non clear change N/A = not assessed

### Knock-out of CG42795 increases the amount of Rab2 structures in larval fat bodies

In addition to RNAi silencing of the *CG42795* gene, we also investigated its knockout variant *Mi{MIC}CG42795[MI11017] (CG42795[Mi]*). For this, we used a strain in which a Minos (a member of the Tc1/mariner superfamily) transposon-based insertion was introduced [43]. The construct, MiMIC (Minos-mediated Integration Cassette), is integrated into the intron of the gene. It contains a splice acceptor, a STOP sequence, reporter genes and a polyA signal. MiMIC disrupts normal splicing, and the extra STOP codon stops transcription, thus resulting in either complete gene knockout or a truncated protein [44]. We isolated total RNA from *CG42795[Mi]* strain and analyzed the amount of *CG42795* mRNA level using PCR. The result showed that the amount of CG42795 mRNA was reduced in *CG42795[Mi]* animals ***(Sup. Fig.: 1)***. We examined the amount of membrane-bound active Rab2 protein in fat bodies using endogenous GFP-Rab2 reporter [45]. In well-fed animals, both RNAi and gene knockout significantly increased the amount of active Rab2, which is also clearly visible in the increase in the number of Rab2-bound vesicles appearing in fat bodies ***(Figure 6: A and Sup. Fig. 2: A)***. When the samples were examined with Lyso-TrackerRed staining, the Minos construct-containing knockout strain significantly increased the amount of acidic vesicles in both fed and starved conditions ***(Figure 6: B)***. CG42795-RNAi caused an increase only in starved animals (***S. Fig.: 2: B***).

**Figure 6:**
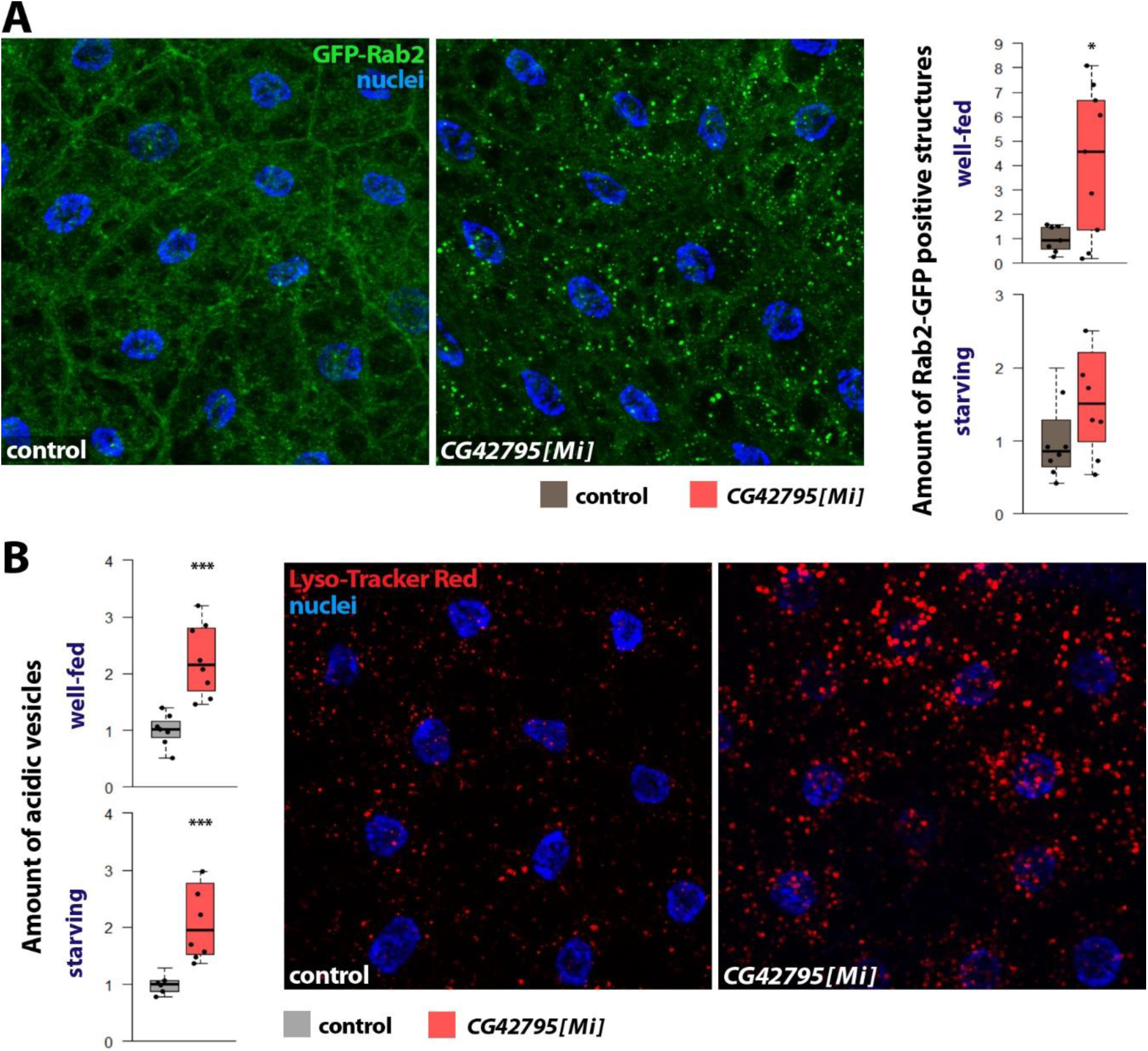
Knock-out of CG42795 increases the amount of active Rab2 and acidic-degrading vesicles. A: The amount of active Rab2 spots increases in the CG42795 knock-out fat bodies in both conditions. **B:** CG42795[Mi] cells show a significant increase of acidic vesicles in both conditions of the Lyso-TrackerRed-stained fat cells. Levels of significance: * = p<0.05; ** = p<0.01; *** = p<0.001.

The human ortholog of the CG42795 gene, TBC1D30 has similar function, regulating Rab2 and autophagic activity.

Taken together the data of our previous experiments, posttranscriptional inhibition of *CG42795* was the most beneficial for activating neuronal autophagy and improving vitality in *Drosophila*. Confident in the conservation of this protein’s functions, we also examined how the silencing of this protein effects human HeLa cells. The human ortholog of the *CG42795* gene is *TBC1D30* (TBC domain family member 30). Experimental studies have not previously investigated the regulatory role of TBC1D30 regarding to Rab2 and autophagy.

For investigating the effect of this conserved gene we silenced the *TBC1D30* gene in the HeLa cells. The efficiency of the RNAi was measured with Q-PCR technique (TaqMan assay) and the results showed that the mRNA product of the gene was knocked down to one tenth of the original amount *(**S. Fig.: 3**)*. For our experiment, we used the RFP-GFP-LC3B fusion protein under confocal microscopy in HeLa cells, a popular tool for the dynamic study of autophagy that allows dual fluorescent labelling [42]. In this fusion protein, the pH stability of RFP is higher that of GFP, so the RFP-GFP-LC3B localized to the inner surface of mature autolysosomes, which have low pH (∼4-5) will show a red signal in confocal microscopy [46]. Following the silencing of the *TBC1D30* gene, the amount of RFP-LC3B-positive vesicles in the cells increased, indicating elevated autophagy activity ***(Figure 7: A and S. Fig. 4: A)***. We did not observe an accumulation of yellow marking, so *TBC1D30-*RNAi did not cause a fusion problem. We examined whether the autophagic flux was functioning properly using Western blot, preparing protein samples isolated from the *TBC1D30-*RNAi human cells containing the RFP-GFP-LC3 marker. We used anti-GFP to label the protein sample, which will produce 4 bands on the Western blot image. The two bands on the top, around 70 kDa are the soluble, cytosolic form (I) and the lipidated form (II) of the intact fusion protein. This intact RFP-GFP-LC3B protein is not yet reached the lysosomal degradation phase. The band at ∼54 kDa contains RFP-GFP fusion without the LC3B. During autolysosome degradation, the LC3B on the C-terminal part cleaves off, while the fluorophores (RFP+GFP) remain intact. The band at ∼27 kDa contains the free fluorophores ***(Figure 7: B)***. During autophagic-lysosomal degradation, the whole fusion protein is separated into free units, but the GFP and RFP parts are more resistant to lysosomal proteases, so they released and visible as separate bands on Western blot therefore making them good indicators of active autophagic flux [46]. Our experiment showed that the autophagic flux is increased furthermore we observed greater autophagic activity, which continues at least until autolysosome formation. In fact, in the *TBC1D30*-RNAi cell samples, we find a higher proportion of RFP-GFP fusion and free GFP proteins, indicated by the stronger middle and bottom bands. The RFP-GFP-LC3B-II bend was also stronger in the RNAi-cells that could mean more autophagic vesicles present ***(Figure 7: B)***. In the next experiment, we measured the amount of active Rab2 protein in GAP-silenced HeLa cells to determine whether there was a correlation with Rab2 regulation. Using confocal microscopy, we found that the amount of structures containing active Rab2 protein had greatly increased in *TBC1D30*-silenced human cells ***(Figure 7: C and S.Fig. 4: B)***. These results indeed suggest a conserved function of the TBC1D30 in human cells and that it could regulate Rab2 protein and autophagic activity.

**Figure 7:**
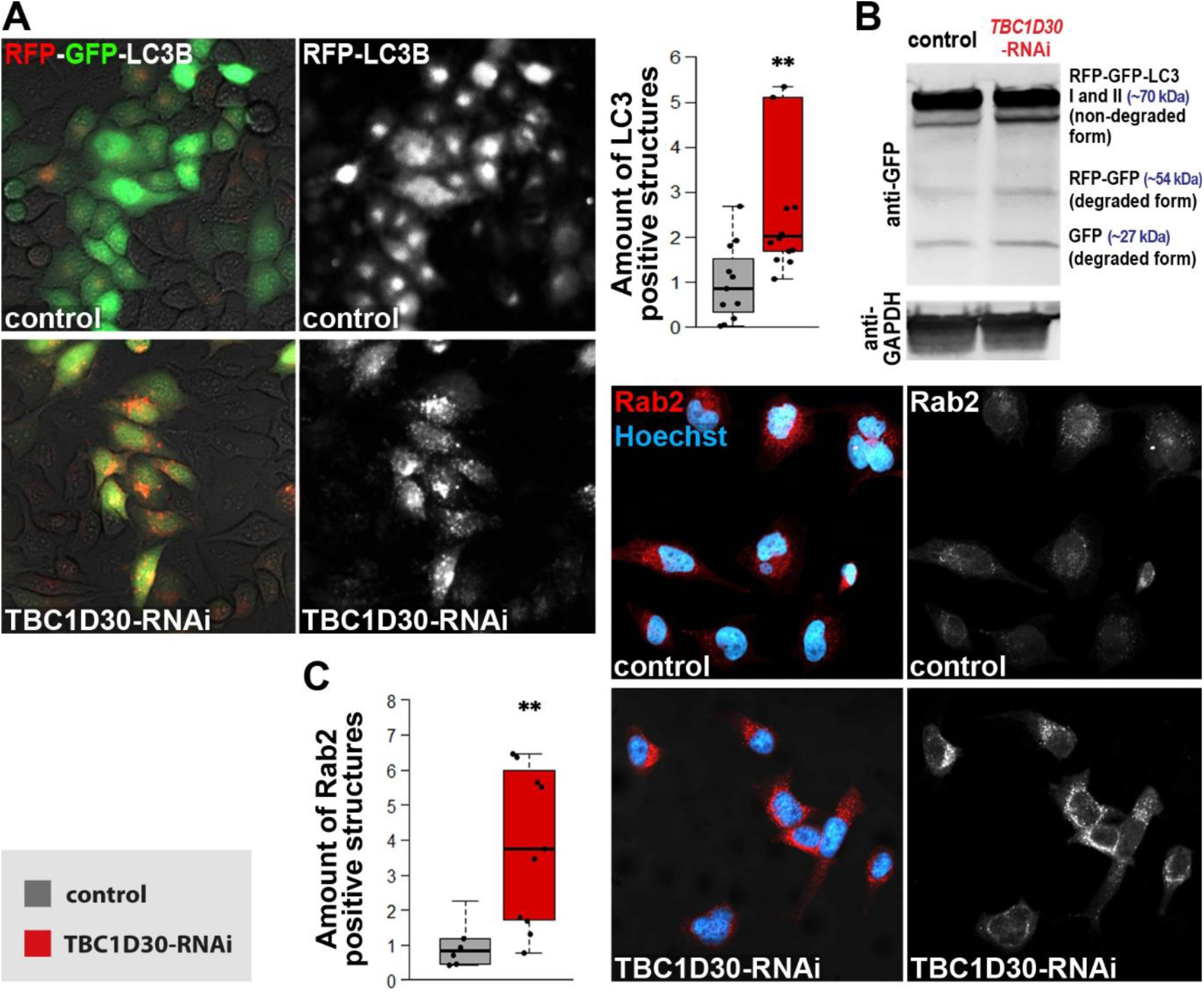
The absence of TBC1D30 increases Rab2 and autophagy activity in HeLa cells. A: Confocal microscopic images of human cells expressing RFP-GFP-LC3B autophagic marker. TBC1D30-RNAi images show more LC3B positive vesicles than the control images. Evaluated data on the right in the form of boxplots. **B:** Western blot image isolated from the modified human cells used in the RFP-GFP-LC3B experiment. An increase of the degraded form of the marker can be seen in the TBC1D30 silenced cells (right column) suggests an increased autophagic activity. **C:** RFP fused Rab2 protein expressing HeLa cells. An increase of active Rab2 protein was detected in the TBC1D30 silenced cells compared to control in confocal microscopy. Levels of significance: * = p<0.05; ** = p<0.01; *** = p<0.001.

To verify whether the human TBC1D30 protein directly interacts with the Rab2 protein, we performed a colocalization experiment. In the experiment, anti-Rab2 and anti-TBC1D30 antibody labeling were used in HeLa cells. The doubly labeled cells were examined with a confocal microscope. The yellow (doubly labeled) fluorescent dots indicate the two proteins are located close to each other ***(Figure 8: A)***. During our experiments, a moderately strong correlation was observed, with the points fitting well along a line in the Pearson correlation test. Furthermore, a strong correlation can be observed in the intensity changes of the two protein signals. The r value of the Pearson coefficient is around 7.5.

**Figure 8:**
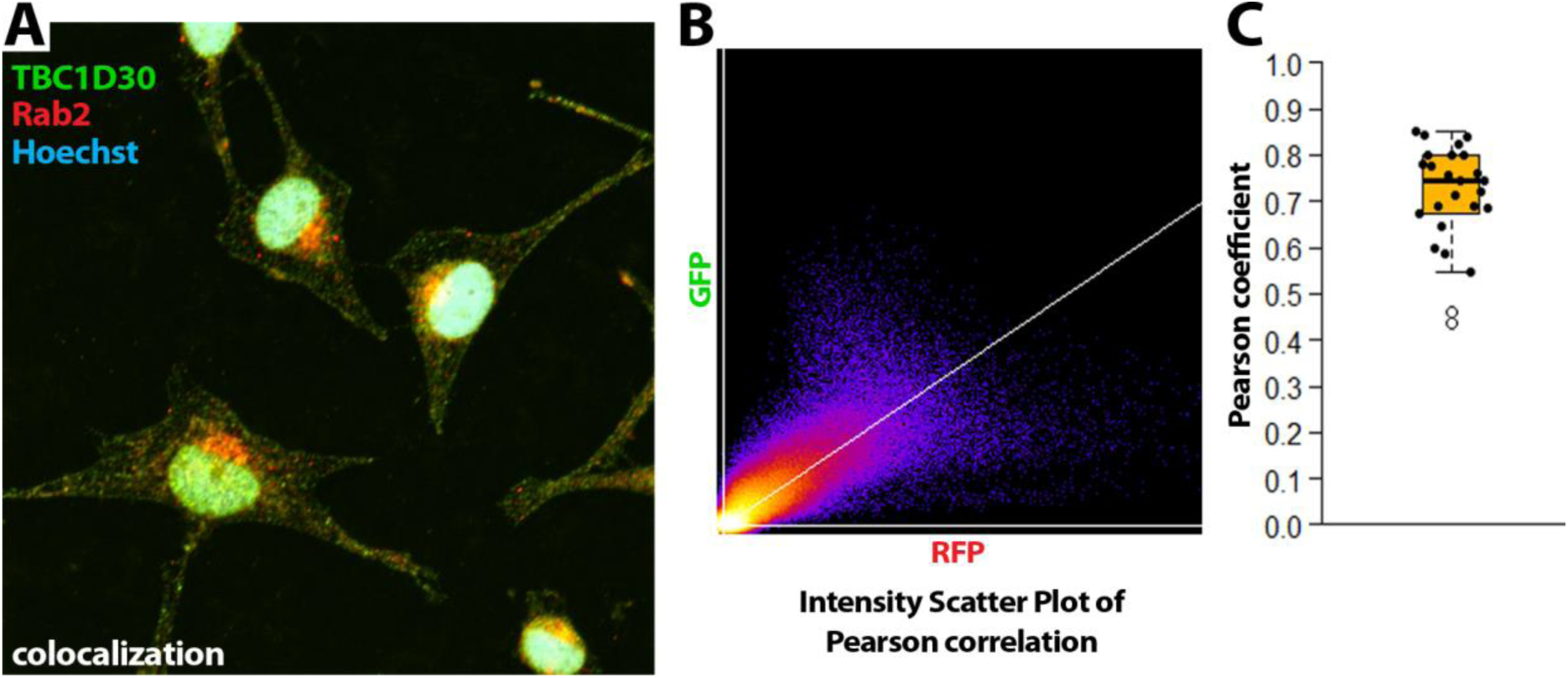
Rab2 and TBC1D30 proteins partially colocalize with each other in HeLa cells. A: Confocal microscopic images of HeLa cells labeled with anti-Rab2 (red) and anti-TBC1D30 (green) antibodies. The nuclei were dyed with Hoechst (blue). As the two proteins making a close interaction the mixture of fluorescent lights turn to yellow spots. **B:** Intensity scatter plot of Pearson correlation showing moderately strong correlation of the colocalization of the two proteins. The colour code shows the pixel density (yellow/red = many pixels, blue/purple = few). **C:** Pearson coefficient plot, where 1 means perfect linear correlation and 0 means there is no correlation between the changes of the intensity of two signals. Each dot on the plot represents separate samples analyzed.

## Discussions

In the present study, we investigated the role of 12 GTPase-activating proteins (GAPs) belonging to the TBC1 domain family in Rab2 and autophagy regulation. Silencing of several GAPs was found to affect the amount of membrane-bound form of Rab2 ***(Figure 2 and Figure 3)***. The role of GAPs was investigated in larval fat bodies and adult animals’ nervous system. Silencing of *CG1695*, *CG11727*, and *CG31935* genes in both organs resulted in Rab2 activation. However, in several cases, differences were found between GAP silencing in larval fat bodies and in adult brains. *CG5344*-RNAi and *CG7324*-RNAi only increased the number of Rab2-positive structures in the nervous system. In contrast, decreased expression of *CG6182*, *CG8085*, *CG42612* altered the amount of YFP-Rab2 reporter only in fat bodies.

After the silencing of GAPs, it was found that in the nervous system, there is no significant difference between Rab2 positive vesicles measured in the cell body and axonal sections ***(Figure 3)***. Alone, *CG7324*-RNAi caused a considerable increase in Rab2 only in the axonal part, but the enhancing effect also showed a tendency to increase in the nuclear part of the neurons. However, we observed significant Rab2 regulatory differences in GAP silencing in fat bodies between fed and starved conditions ***(Figure 2)***. *CG6185* and *CG42612* increased the amount of Rab2-positive structures in well-fed samples only. In contrast, silencing of *CG4041*, *CG11727*, *CG12241*, and *CG42795* genes increased Rab2 activity only in starved samples (lysosome amount was significantly increased by *CG42612*, *CG11727*, and *CG31935* silencing in both conditions tested). Similar condition-dependent differences were also observed for other autophagy regulators. The *Drosophila* orthologue of myotubularin-type phosphatase 14 (MTMR14) inhibits basal autophagy exclusively in fat bodies, whereas Mtmr6 (common orthologue of human MTMR6,-7,-8) is a specific inhibitor of stress-induced autophagy [47]. Comparing the two examined organs, the roles of two GAPs, *CG11727* and *CG31935*, seem to be common. While, for example, *CG42795* is only able to play its role in the nervous system. According to the Fly Cell Atlas, *CG42795* is expressed at lower levels in larval tissues, and in adult animals it is most intensively expressed in the nervous system (inter-, motor-, sensory-neurons and glia cells). This may explain why its silencing had a phenotypic result mainly in the nervous system. The expression of *CG11727* is high in the adult nervous system and in L3 larvae [48], [49]. In *Drosophila*, Rab2 is known to regulate the fusion of various types of vesicles with lysosomes and their transport in different organs. In fat bodies, Rab2 regulates the fusion of Golgi derived vesicles with lysosomes but does not affect autophagosome - lysosome fusion [50]. In contrast, in larval neurons, eye primordia and muscle, Rab2 promotes autophagosome-lysosome fusion [51], [52]. When we examined autophagic processes in the brains of fruit flies, silencing the *CG42795* gene resulted in significant changes in the levels of several autophagy substrate proteins. Both Atg8a and Ref(2)P were reduced, and the presence of ubiquitin-tagged structures was also significantly reduced in the flies’ neurons. *CG1695*-RNAi also showed similar results in these tests *(Figure 4)*. These may indicate more efficient acidic degradation of the mentioned autophagic substrate proteins. We investigated the effects of 10 selected GAPs silencing on the animals’ lifespan and locomotion ability. *CG8085*-RNAi and *CG42795*-RNAi extended the lifespan in both sexes, whereas *CG7324*-RNAi and *CG12241*-RNAi had a positive effect only in males ***(Figure 5)***. The silencing of *CG42795* and *CG7324* positively affected the animals’ locomotion ability, compared to *CG8085*-RNAi, which impaired locomotion ability.

Taken together our data, posttranscriptional inhibition of *CG42795* was the most beneficial for activating neuronal autophagy and enhance vitality in animals. Previously *CG42795* was only described as a GTPase activating protein which main role can be a Rab-GAP [FlyBase, FBgn0261928; NCBI Gene ID: 41252], [53]. However, in a recent study this protein was also found in the presynaptic active zones (AZ) of fruit fly neurons. The researchers found that a specific isoform of CG42795 is able to associate with the scaffolding protein Bruchpilot (BRP) and is required for the formation of proper presynaptic AZs. Their experiments revealed that a lack of *CG42795* in the mushroom body impaired olfactory aversive memory consolidation in flies and resulted in ectopic AZ aggregates (“blobs”), a phenotype after which the gene was named *Blobby* [54]. In our experiments, we investigated the role of the *CG42795/blobby* in terms of autophagy and Rab2 protein. Since silencing this GAP performed the best among those tested therefore, we investigated the role of the human orthologue, TBC1D30. Based on the data we have available TBC1D30 working as a Rab-GAP protein negatively regulating Rab8 [55], [56], [57] and it also plays a role in the repression of proinsulin and insulin secretion from the cells of human Langerhans-islands [58], [59], [60]. TBC1D30 also could be a negative regulator of Rab3 smallGTPase [58]. Rab8 is responsible for transport between the trans-Golgi network and the plasma membrane, while Rab3 regulates synaptic vesicle storage and neurotransmitter release in synapses [61], [62]. These findings came from mammalian experiments, but the study of the corresponding Drosophila orthologs may also be useful, which could serve as the basis for our next research topic. Anyways, in our experiments knocking down TBC1D30 caused an increase in the number of active, membrane bound form of Rab2 protein and enhanced the activity of autophagy without blocking autophagic flux. TBC1D30 also showed moderately strong colocalisation with the Rab2 protein. Since GAPs only perform a GTPase activating function by interacting with small GTPase proteins, which does not require long-term close association, it cannot be expected that the two proteins are in a continuous close interaction with each other in a large percentage of cases. A GAP and a Rab protein remain together until the GTP-GDP conversion occurs in the Rab protein, after which the GAP leaves the Rab protein [63], [64]. Exact data is not available, and the binding time may vary between Rab-GAP pairs, but the catalysis reaction time takes approximately 1 second [65]. These findings suggest that this protein may play a significant role in regulating the autophagic pathway and the Rab2 protein in human cells. Furthermore, since we experienced similar effects when we silenced the *Drosophila* ortholog of TBC1D30 we can assume that this gene has a conserved function in humans. Proximity labelling and yeast two-hybrid experiments have identified three additional GAP genes that may interact with Rab2 in mammalian cells: TBC1D9, TBC1D10B and TBC1D25 [66], [67], [68]. Investigating these genes may reveal important regulatory points. Further study of *CG42795* and TBC1D30 may also help to understand the regulation of autophagy and Rab-specific small GTPases. Additional experiments may reveal a new potent lifespan and autophagy regulatory point.

The *Drosophila* TBC1 domain family includes 25 proteins, of which we have investigated the roles of 12. It might be worthwhile in the future to investigate the regulatory role of the additional 13 GAPs on Rab2 and autophagy. In our previous research, we have also investigated the effects of the neuronal activation of Rab7 and Arl8. Arl8 was found to have a positive effect on neuronal activation similar to Rab2 activation. Autophagy was activated in animals, locomotor function improved, and lifespan extended. In contrast, overexpression of the constitutively active form of Rab7 in dopaminergic and serotonergic neurons had the opposite effect to the activation of Rab2 and Arl8 [25]. In the future, it might be worth investigating the regulatory points of Arl8 and Rab7 similarly to our current study and finding regulatory junctions that can co-activate Arl8a and Rab2 without affecting Rab7 activity.

Our results have highlighted that multiple GAPs have the capacity to regulate a single small GTPase. Similarly, a single GAP has the capacity to regulate a multitude of distinct small GTPases. Consequently, future research should prioritize studying the functions of the investigated GAPs on other Rab enzymes and to explore the compensatory mechanisms among GAPs with analogous regulatory functions.

## Author Contributions

G.F. and F.K. performed experiments on *Drosophila* samples, analysed data and wrote the manuscript. T.M.B. and Gy.Sz. performed experiments on *Drosophila* samples and analysed data. O.K., Zs.S-V. completed experiments on human cells. Sz.T., P.L. and O.T. wrote and edited the manuscript and provided financial support for the work.T.O. and P.L. performed experiments on human cells, analysed data and wrote the manuscript. T.K. designed experiments, analysed data, and wrote the manuscript and provided financial support for the work.

## Acknowledgments

*Drosophila* strains and reagents were kindly provided by Gábor Juhász and Ole Kjaerulff. The authors also thank Regina Preisinger, Beatrix Supauer, Erzsébet Gatyás for the excellent technical assistance.

