## Supplementary material for "The GTPase-activating protein CG42795 is a potent neuronal regulator of ageing in *Drosophila melanogaster*": S.Figures: Falcsik _et_al._Supplementary_2026.pdf

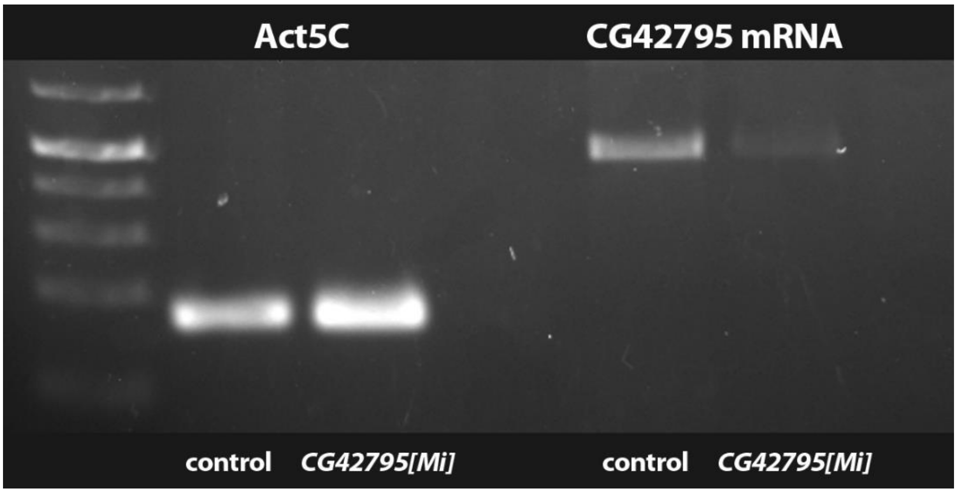


***Sup. Fig. 1: CG42795[Mi] significantly reduced the amount of CG42795 mRNA in fruit flies.*** *KO validation by RT-PCR. Left: bands of Actin5C housekeeping control mRNA in control and CG42795[Mi] flies. Right: amount of CG42795 mRNA.*

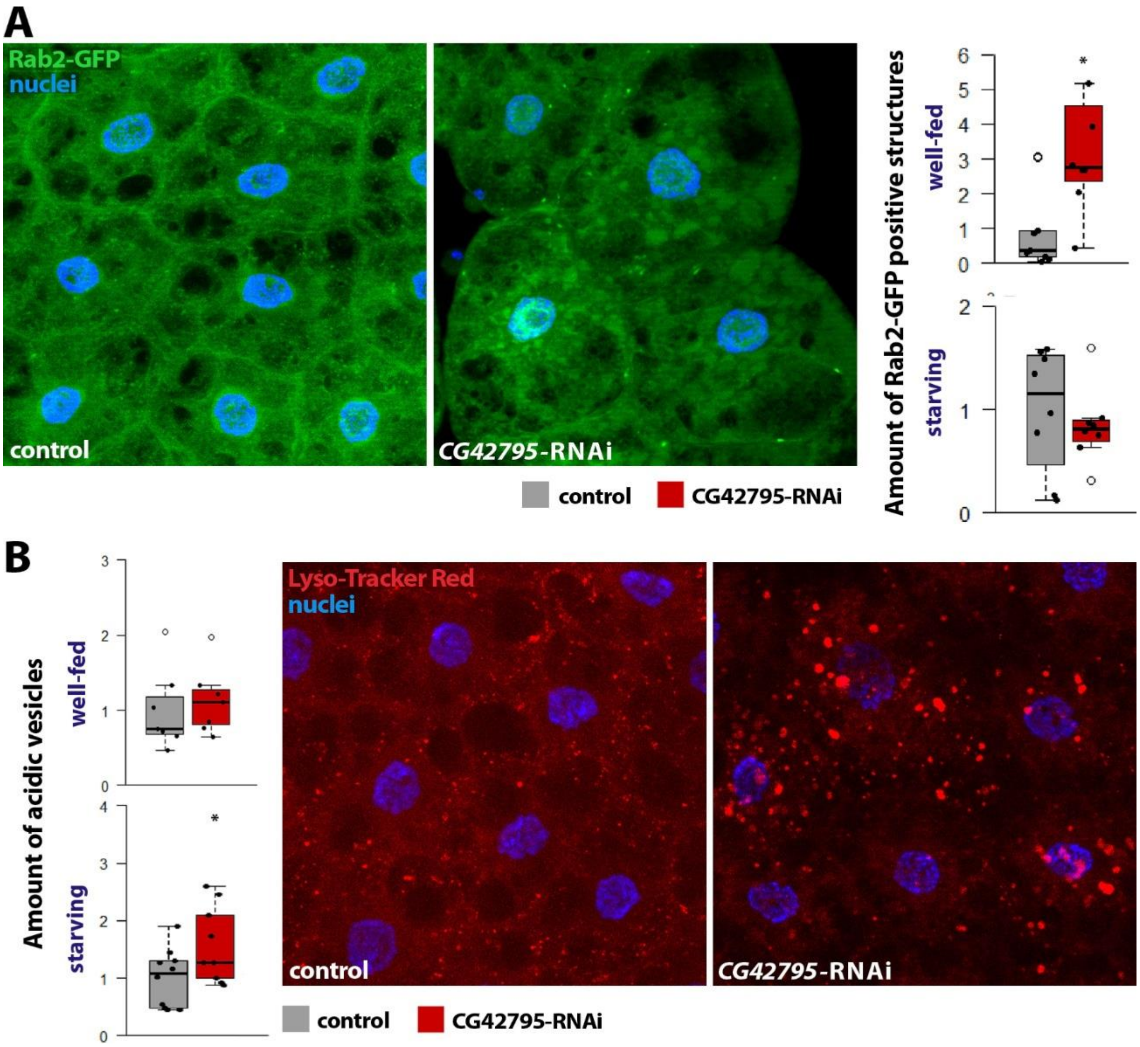


***Sup. Fig. 2: RNAi of CG42795 increases the amount of active Rab2 positive structures and acidic vesicles in Drosophila fat bodies. A:*** *YFP-Rab2 positive structures appear in the CG42795-RNAi fat bodies.* ***B:*** *Lyso-tracker Red staining of the CG42795-RNAi fat bodies shows that the amount of acidic*

*vesicles increased in these cells compared to the control in starving condition.* ***Level of significance: * = p<0.05***

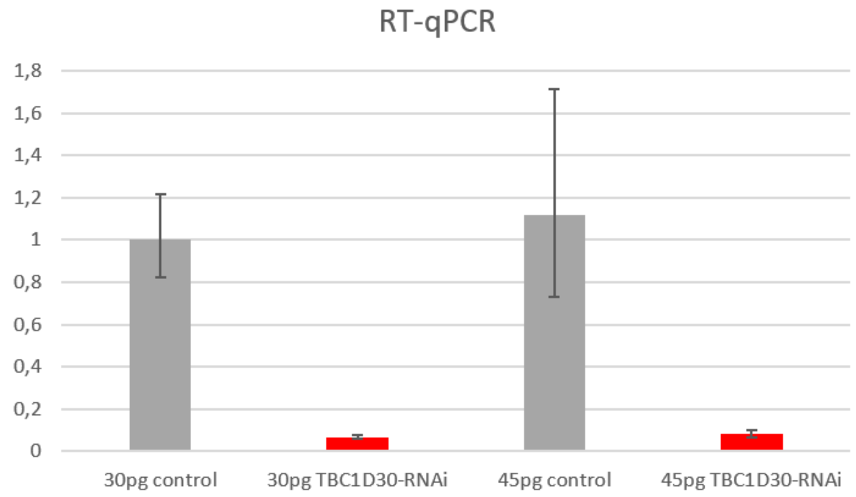


***Sup. Fig. 3: Silencing TBC1D30 in HeLa cells significantly reduced TBC1D30 mRNA levels.*** *qPCR data of control and TBC1D30 silenced HeLa cell RNA isolate. RNA isolated from 30pg sample (left) and 45pg sample (right).*

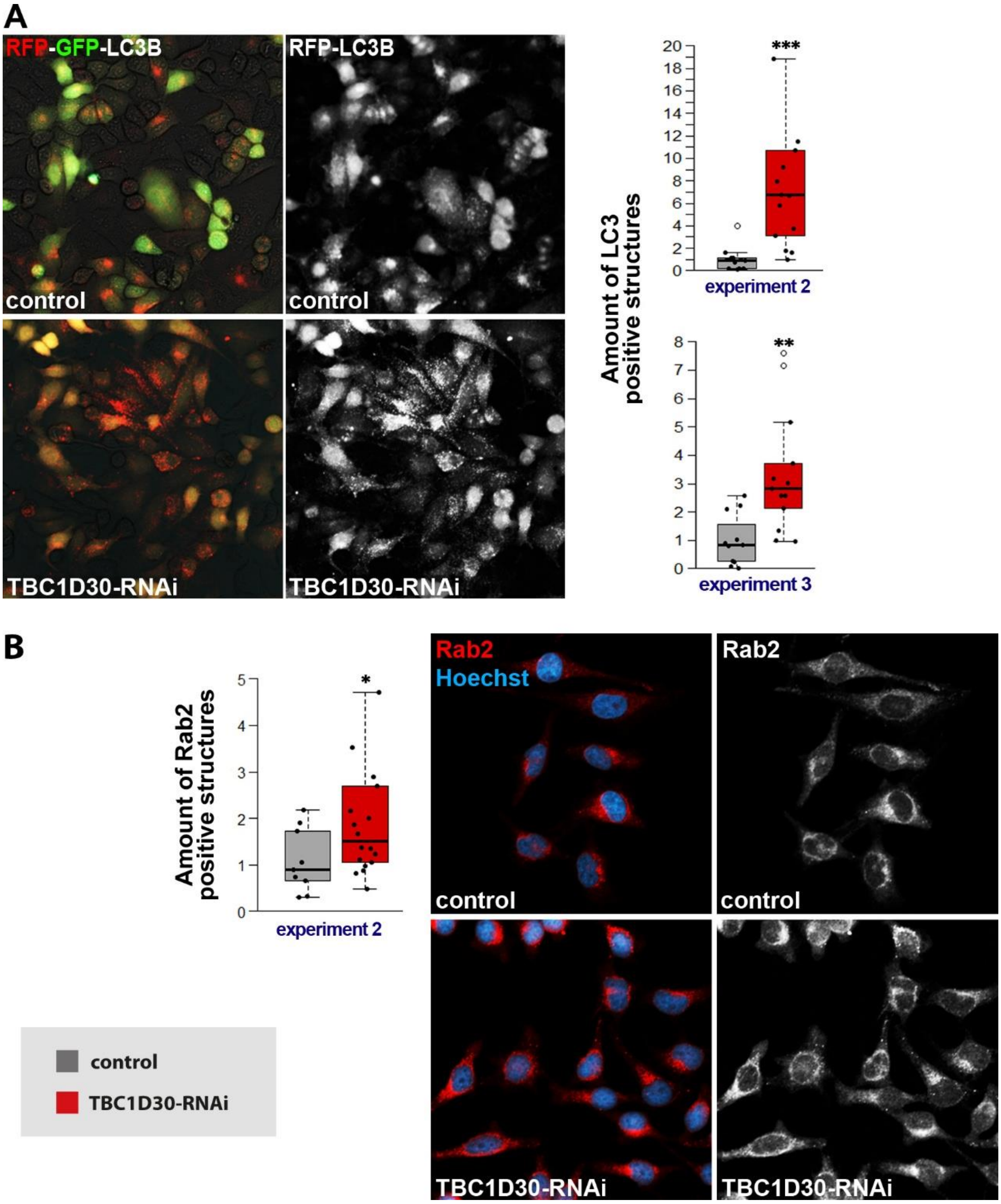


***Sup. Fig. 4: TBC1D30 silenced HeLa cells shows an increase in autophagic vesicles and active Rab2 positive structures. A:*** *RFP-GFP-LC3B proteins emits RFP-LC3 in acidic environment (autolysosomes), the TBC1D30-RNAi cells increases the amount of LC3-positive structures, indicating enhanced autophagy.* ***B:*** *Immune labeling of Rab2 protein in HeLa cells. The amount of active Rab2 positive (membrane bounded) structures increase in the TBC1D30-RNAi cells.* ***Levels of significance: * = p<0.05; ** = p<0.01; *** = p<0.001***
